# ATAC-seq identifies thousands of extrachromosomal circular DNA in cancers and cell lines

**DOI:** 10.1101/845040

**Authors:** Pankaj Kumar, Shashi Kiran, Shekhar Saha, Zhangli Su, Teressa Paulsen, Ajay Chatrath, Anindya Dutta

**Author notes:** Corresponding Author: Anindya Dutta.

## Abstract

Extrachromosomal circular DNAs (eccDNAs) lack centromeres and are not passed equally into daughter cells and thus provide a source of cell-cell heterogeneity in normal and tumor cells. Previously we and other groups purified circular DNA and linearized them by rolling circle amplification for paired end high throughput sequencing to identify eccDNA in cells and tissues. We hypothesized that many eccDNA will have more open chromatin and so will be susceptible to linearization and sequencing by tagmentation. Indeed, we find that ATAC-seq on cell lines and cancers, without any enrichment of circular DNA, identifies thousands of eccDNAs. The identified eccDNAs in cell lines were validated by inverse PCR on DNA that survives exonuclease digestion of linear DNA and by metaphase FISH. We demonstrate that ATAC-seq data generated in Lower Grade Gliomas (LGG) and Glioblastomas (GBM) identify eccDNAs, including one containing the well-known EGFR gene amplicon from chr7. Many of the eccDNAs are identified even before amplification of the locus is detected by genotyping arrays. Thus, standard ATAC-seq is a sensitive method to detect segments of DNA that are present as eccDNA in a subset of the tumor cells, ready to be further amplified under appropriate selection, as during therapy.

## Introduction

**ATAC-seq** (**A**ssay for **T**ransposase-**A**ccessible **C**hromatin using **seq**uencing) identifies open chromatin regions all across the genome (Buenrostro et al., 2015). The method uses the hyperactive transposase Tn5 to cut the accessible chromatin with simultaneous ligation of adapters at cut sites (Buenrostro et al., 2015). To reduce the contamination of mitochondrial DNA in library preparation the nuclei pellets are isolated first from cells or tissues before tagmentation step (Corces et al., 2018).

We previously reported the presence of tens of thousands of extrachomosomal DNA (eccDNA) in the nuclei of human and mouse cell lines as well as normal tissues using paired end high throughput sequencing of circular DNA enriched preparations (Dillon et al., 2015; Kumar et al., 2017; Shibata et al., 2012). Several other groups have also used similar approaches to describe the presence of eccDNAs in various eukaryotes ranging from yeasts to humans (deCarvalho et al., 2018; Moller et al., 2015a; Moller et al., 2018; Moller et al., 2015b; Shoura et al., 2017; Turner et al., 2017). Since isolated nuclei as a whole are subjected to the transposition reaction in ATAC-seq, we hypothesized that the transposase will also cleave DNA from eccDNAs, and so the ATAC-seq libraries will contain fragments of DNA from eccDNA.

To test our hypothesis we first prepared ATAC-seq libraries using C4-2B (prostate cancer) and OVCAR8 (ovarian cancer) cell lines and identified hundreds of eccDNAs using our newly developed computational pipeline. Inverse PCR on exonuclease resistant extrachromosmal DNA (highly enriched in circular DNA) and FISH on metaphase spreads confirmed the presence of the identified somatically mosaic eccDNA. To provide additional evidence of the success of ATAC-seq in identifying eccDNA, we analyzed an ATAC-seq library generated from patient derived GBM cell lines (Xie et al., 2018) and identified the eccDNA harboring EGFR gene, which is known to be amplified through the formation of eccDNA in GBM. Finally, we analyzed ATAC-seq data from GBM and LGG generated by the TCGA consortium to identify hundreds of eccDNAs even before their amplification in the tumor was apparent as a copy number variation by hybridization to SNP arrays. Genes involved in pathways related to nucleosomal events were significantly enriched in these loci.

## Results

### Principle of circular DNA identification by tagmentation method

EccDNAs are known to have chromosomal origin. A linear DNA fragment is generated either by the chromosome breakage due to adjoining DNA breaks, e.g. in chromothripsis (Maher and Wilson, 2012), or by DNA synthesis related to DNA replication or repair. The two ends of a linear DNA are ligated to make a circular DNA (Figure 1A), creating a specific junctional sequence that is not present in the normal reference genome. We have developed a very simple method to identify eccDNAs by collecting all the read pairs where one read of a pair maps uniquely to the genome in a contiguous manner (<=5 bp insertions or deletions or substitutions) and the other read maps as a split read (non-contiguous segments that could be as far apart as few MB, but usually are much closer) flanking the mapped read (Figure 1B). The split read maps to the circular DNA ligation junction and the other (contiguously mapped read) maps to the body of the putative eccDNA. The start of the 1^st^ split read and the end of 2^nd^ read are annotated as start and end of eccDNA. Tandem duplication of DNA in the genome will also create a similar junctional sequence, but for the purpose of identifying incipient gene amplification, an eccDNA or a tandem duplication of a chromosomal segment are equally important. However, if the aim is to exclusively and comprehensively identify eccDNA the ATAC-seq library should be prepared from eccDNA enriched samples where linear DNA has been digested with exonucleases. The complete pipeline to identify eccDNA coming from one locus (non chimeric eccDNA) of any length is available through our GitHub page (https://github.com/pk7zuva/Circle_finder (https://github.com/pk7zuva/Circle_finder/blob/master/circle_finder-pipeline-bwa-mem-samblaster.sh)). The steps to find a circular DNA from any paired end high throughput-sequencing library are detailed in Figure 1B-C and Methods.

**Figure 1:**
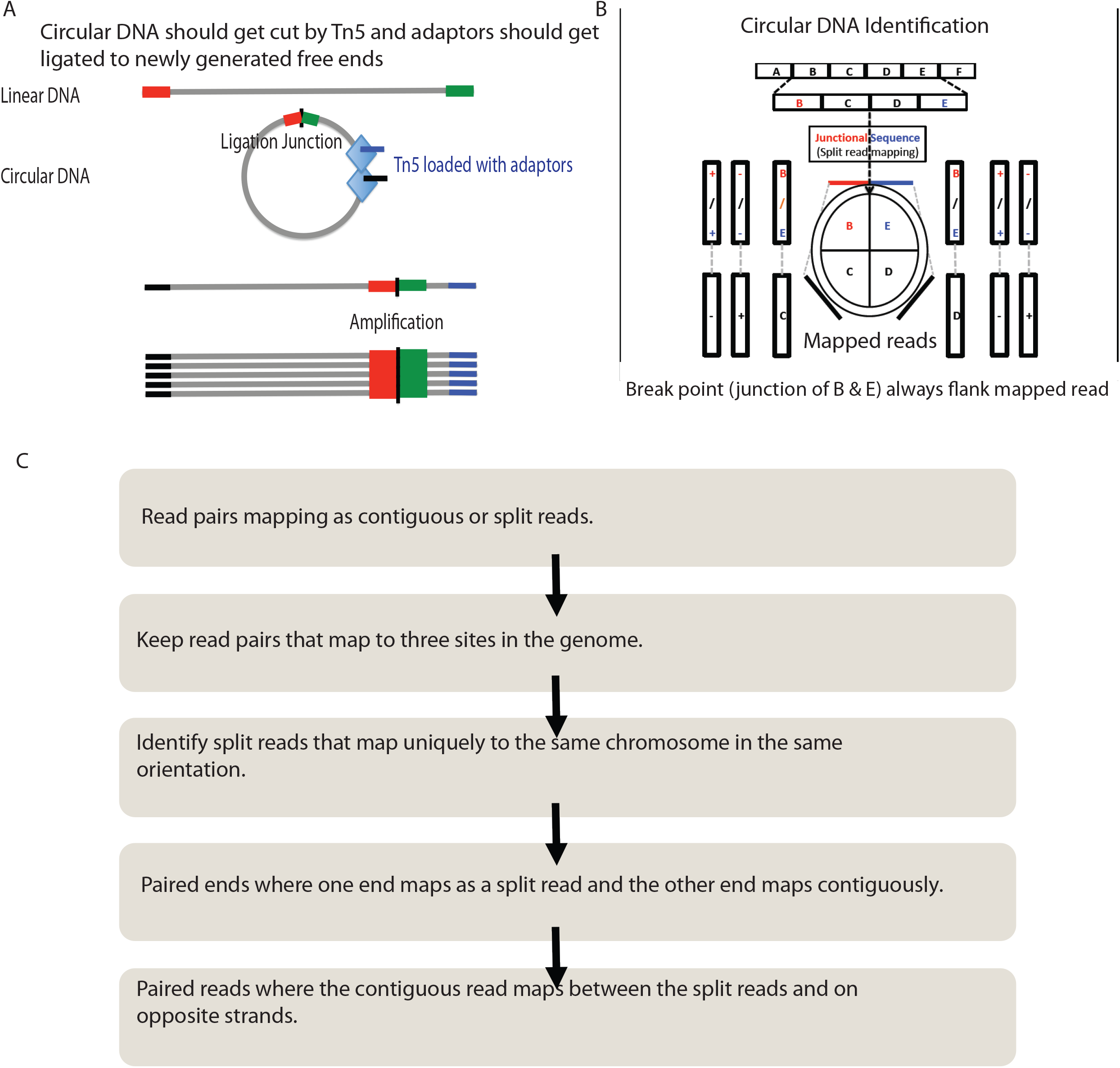
A schematic to show that circle could be part of an ATAC-seq library. **(A)** If circular DNA has open chromatin structure near or around the ligation point the library preparation method will cut and attach an adaptor into a DNA fragment from eccDNA. **(B)** shows one end of paired end read mapping on the body of a circular DNA with read from the other end mapping on the ligation junction. **(C)** shows detailed steps from mapping to identification of the new Circle_finder pipeline.

### Application of ATAC-seq method to identify circular DNA in OVCAR8 and C4-2B cell lines

We prepared ATAC-seq libraries from C4-2B prostate cancer and OVCAR8 ovarian cancer cell lines. The sequencing and mapping statistics are given in Table 1. >90% of the reads mapped to human genome and the computational pipeline identified hundreds of circular DNA. The length distribution of eccDNA is shown in Figure 2A: around 64% in C4-2B and 43% in OVCAR8 of eccDNA are <1 kb, and so are similar to the microDNAs that we identified earlier in normal and cancer cells (Kumar et al., 2017; Shibata et al., 2012). However, 36% of the eccDNA in C4-2 and 57% in OVCAR8 are >1 kb, including eccDNAs long enough to encode gene segments or even complete genes. The eccDNA are derived from all the chromosomes (Figure 2B). As a positive control we identified hundreds of junctional sequences from the circular mitochondrial genome (data not shown).

**Table 1:** Summary of eccDNA sequencing and mapping to the human genome in C4-2 & OVCAR8 cell lines.

|  | C4-2 | OVCAR8 |
| --- | --- | --- |
| Total PE150 reads | 374,931,197 | 346,833,854 |
| Mapped in pairs | 372,561,742 | 345,127,010 |
| Mappred-Reads | 373,585,003 | 343,541,464 |
| Un-Mapped-Reads | 1346194 | 1,706,844 |
| #eccDNA | 195 | 171 |

**Figure 2:**
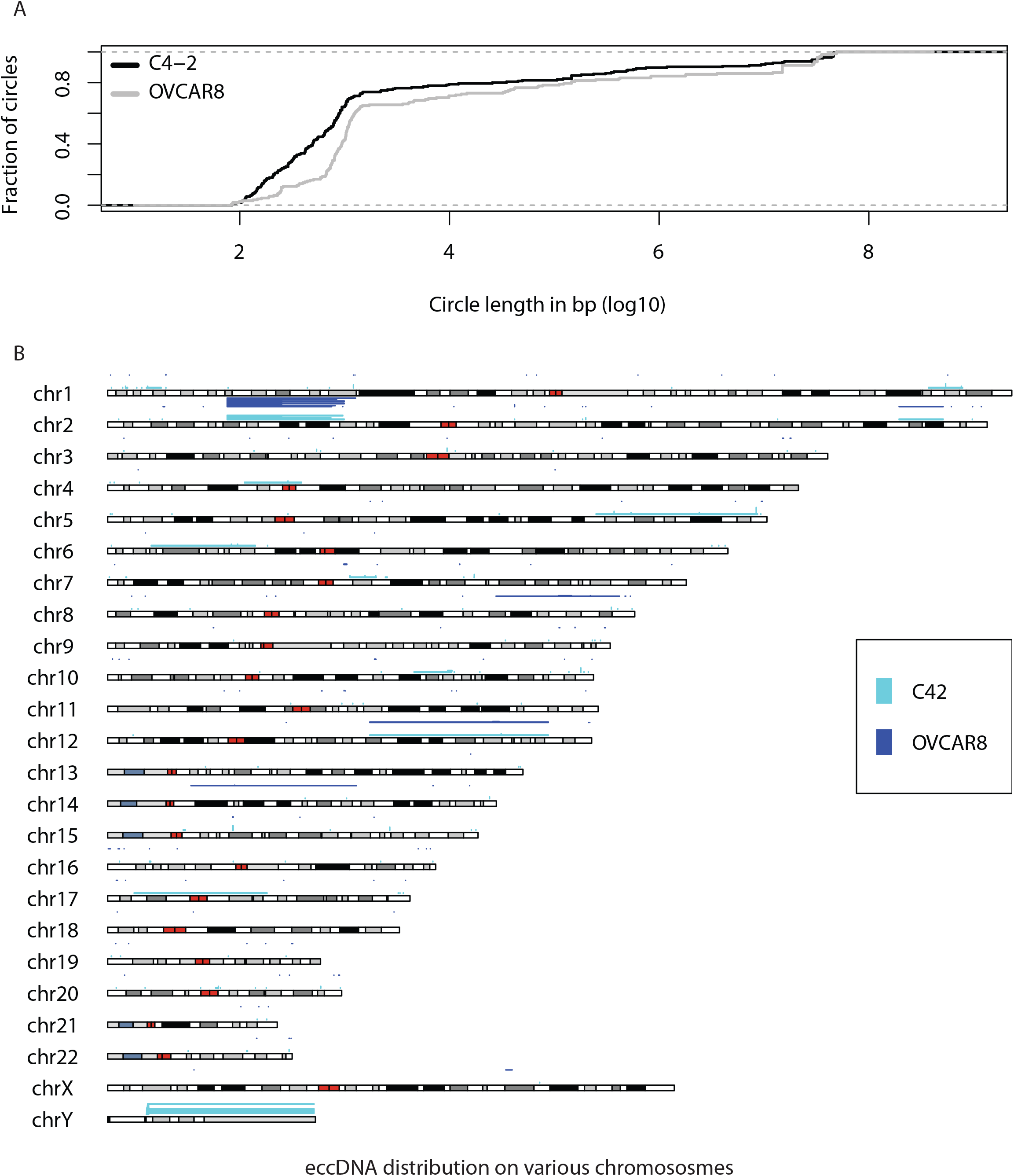
eccDNA in C4-2 and OVCAR8 cell lines. **(A)** Length distribution of identified eccDNA in C4-2 and OVCAR8 cell lines. **(B)** Karyotype plot showing chromosomal distribution of C4-2 and OVCAR8 cell lines.

### Validation of eccDNA identified in C4-2 and OVCAR8 cell lines by inverse PCR

To confirm that the identified junctions are genuinely from eccDNA, and not from tandem genome duplications, we isolated circular DNA by our previously described method that relies on column chromatography and exonuclease digestion to remove all linear DNA and enrich eccDNA (Figure 3A (Shibata et al., 2012)). Inverse PCR was performed with primers designed to amplify across the junctions of eccDNAs from C4-2B & OVCAR8 cell lines (Figure 3B). 11 eccDNAs from OVCAR8 and 6 from C4-2B were tested. 9 of the 11 targets from OVCAR8 and 2 of the 6 from C4-2B gave amplicons of expected sizes (Figure 3B-C). Sanger sequencing of the amplicons confirmed the Junctional sequences identified by ATAC-seq (Figure 3D). A fraction of the primers (2 in OVCAR8 and 4 in C4-2B) did not give desired amplicons possibly because of their presence in low complexity regions or because they came from tandem linear chromosomal duplications which did not survive column chromatography and exonuclease digestion.

**Figure 3:**
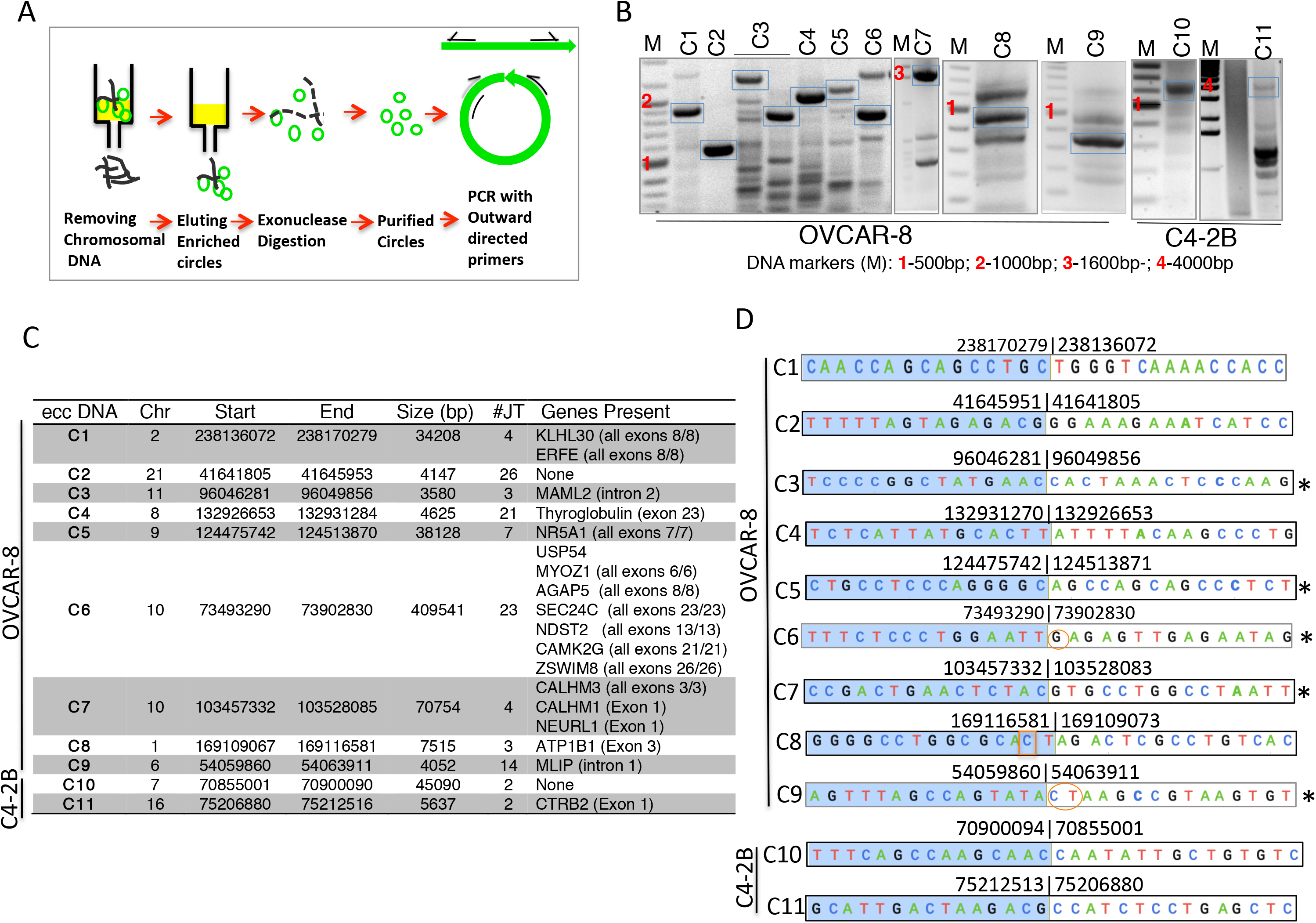
Experimental validation of randomly selected eccDNA identified by ATAC-seq in C4-2B and OVCAR8 cells. **(A)** Schematic for isolation and detection of extra-chromosomal circular DNA. See methods for details. **(B)** PCR detection of eccDNA. DNA bands marked with blue boxes were gel purified and sequenced. **(C)** Description of eccDNAs validated in (B), based on analysis of ATAC-seq data from OVCAR8 and C4-2B **(D)** Junctional tags obtained after sequencing of PCR products in (B). Shaded (blue) and un-shaded sequences depict 15 bases on either side of junctions. Numbers indicate chromosomal location on respective chromosomes. Note the match between numbers for each circle in (C) and (D). Some of the junction sequence identified by Sanger sequencing differ by few bases due to multiple species of eccDNA present in given cell lines. Oval circles represent insertion and boxed sequences represent mismatches. *-Sequence obtained from the bottom strand.

### Metaphase FISH

An independent method for ascertaining whether a locus identified in this study is in an extrachromsomal DNA is to carry out FISH on metaphase spreads. We performed this analysis with two loci that were predicted to be present as either an eccDNA or a gene duplication in OVCAR8 cells: chr2: 238136071-238170279 and chr10: 103457331-103528085. Signal was detected off the main chromosomes in some of the metaphase spreads, but not others (Figure 4A), consistent with the hypothesis that the junctional sequences identify somatically mosaic eccDNA in this cell line.

**Figure 4:**
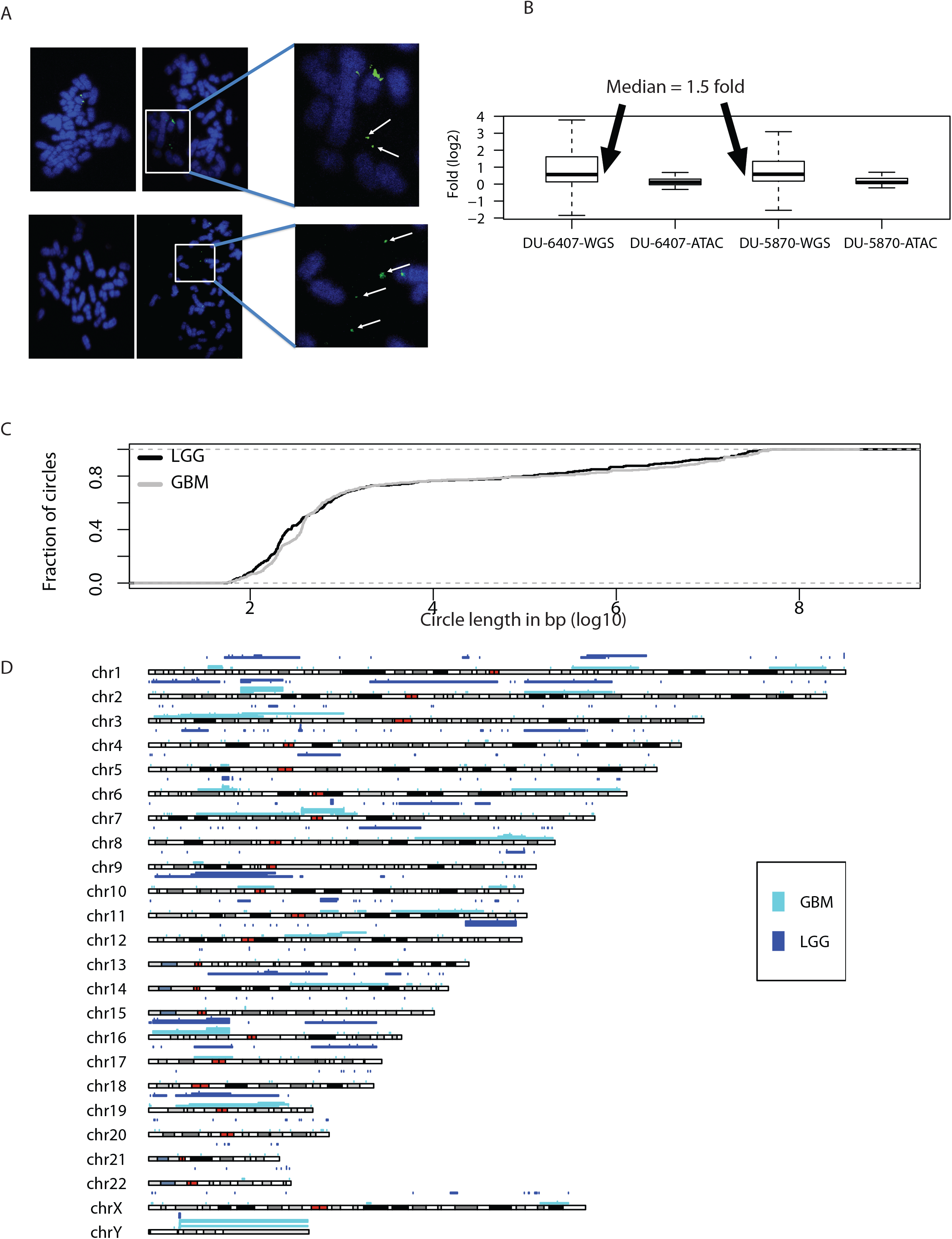
eccDNA in cell lines and LGG or GBM tumors. **(A)** Detection of EccDNA in OVCAR8 cell line by FISH: Metaphase spread of chromosome (blue) from OVCAR8 cells were stained with the probe (green) against the eccDNA locus chr2: 238136071-238170279 (Top Row) or chr10: 103457331-103528085 (Bottom Row). The spreads on the left do not have an extrachromosomal signal, while the spreads on the right have extrachromosomal signals that are better seen in the magnified insets on the extreme right. White arrowheads mark the eccDNA signals. **(B)** eccDNA/duplication loci identified in WGS libraries show genomic amplification (median 1.5 fold) suggesting at least one allele is duplicated in all the cells. eccDNA loci identified in ATAC-seq libraries do not show genomic amplification (median close to zero), suggesting that the eccDNA are apparent before a CNV can be detected at the locus. Value of CNA in Y-axis is in log2. **(C)** Length distribution of identified eccDNA in LGG and GBM TCGA ATAC-seq data. **(D)** Karyotype plot showing chromosomal distribution of LGG and GBM TCGA ATAC-seq data.

### Identification of eccDNA from ATAC-seq data for glioblastoma (GBM) cell lines

Epidermal growth factor receptor (EGFR) was one of the first oncogenes identified in brain cancer and is massively amplified in some GBM patients (Libermann et al., 1985). This somatic copy number variation is present in 43% of GBM patients (Maire and Ligon, 2014). Recent studies have provided further evidence that this oncogenic amplification occurs on eccDNA (deCarvalho et al., 2018; Turner et al., 2017; Xu et al., 2019). To check if we can detect the eccDNA in ATAC-seq data generated from GBM cell lines we turned to six ATAC-seq libraries generated from GBM cell lines developed from a single glioblastoma patient (Xie et al., 2018). We ran the Circle_finder pipeline combining all the six libraries (GSM3318539, GSM3318540, GSM3318541, GSM3318542, GSM3318543 and GSM3318544) and found 58 eccDNAs varying in size from few hundred bases to few megabases. The length distribution and chromosomal distribution of identified eccDNAs are shown in **Supplementary Figure 1A-B**. Most importantly, eccDNA harboring the EGFR gene was the most abundant eccDNA. The top five most-abundant eccDNAs (or tandem gene duplications) identified were chr4:118591708-119454712 (METTL14, SEC24D, SYNPO2, MYOZ2, USP53, C4orf3, FABP2), chr7:54590796-55256528 (SEC61G, EGFR), chr7:54771165-54782815(No protein coding genes), chr7:65038261-65873269 (transcribed unprocessed pseudogenes), chr7:65038264-65873256 (transcribed unprocessed pseudogenes) (**Supplementary Figure 1B**).

### Application to GBM and LGG TCGA ATAC-seq data

Having demonstrated above that ATAC-seq data can be repurposed to identify eccDNA, we turned our attention to ATAC-seq data generated by TCGA consortium (Corces et al., 2018) with a primary focus on two LGGs for which we have whole genome sequencing data and ATAC-seq data. In the TCGA-DU-5870-02A ATAC-seq library we found 21 eccDNAs (junctional tag >=2; 10>1KB, 6>50KB). In the TCGA-DU-5870-02A WGS library we found 749 eccDNAs (junctional tag >=2; 319>1KB, 143>50KB). We further compared the eccDNAs identified in ATAC-seq and WGS libraries and found 25 common eccDNAs (junctional tag>=1; **Supp. Table 1**).

In TCGA-DU-6407-02B ATAC-seq library we found 54 eccDNAs (junctional tag >=2; 15>1KB, 13>50KB) and in TCGA-DU-6407-02B WGS libraries we found 489 eccDNAs (junctional tag >=2; 280>1KB, 136>50KB). 26 common eccDNAs were identified in both libraries (junctional tag>=1; **Supp. Table 2**). Many of common eccDNAs had a high number of junctional tags in the WGS library, perhaps a surrogate marker of their abundance.

**Table 2:** Driver gene amplified on various tumor type

| Tumor Type | Driver gene present on eccDNA |
| --- | --- |
| BLCA | SF1 |
| BRCA | SF1, ERBB2, GNA11, GNA11 |
| COAD | PPP2R1A, MSH3, EEF1A1 |
| ESCA | KRAS, KLF5, ERBB2, KIT, UNCX, MYC |
| GBM | CDK4, EGFR, MYC |
| KIRC | SF1, UNCX |
| KIRP | SF1 |
| LGG | WT1, MACF1, GNA11, KIT |
| LIHC | KRAS, CHD4, GRIN2D |
| LUAD | PPP2R1A, FGFR3, EEF1A1 |
| PRAD | ERBB2, MSH3, UNCX |
| STAD | KRAS, ERBB2, GRIN2D, SF3B1, IDH1 |

We see a higher number of eccDNA/duplication events in WGS compared to ATAC-seq, but 25 and 26 eccDNAs were common (between ATAC-seq and WGS) in TCGA-DU-5870-02A and TCGA-DU-6407-02B libraries respectively (**Supp. Table 1, 2**). The lack of more overlap between the eccDNAs identified by ATAC-seq and WGS from even the same tumor is most likely due to somatic mosaicism (a) because different sections are used for the two libraries and (b) because of insufficient depth of sequencing in either library.

As mentioned earlier, the Circle_finder algorithm cannot distinguish between an extrachromosomal circle and chromosomal segmental tandem duplication without experimentally purifying the circles before library preparation, and so we will refer to these loci as eccDNA/duplication. The signal for the eccDNA/duplication detected from WGS data was strong in the two tumors and was also evident in Copy Number Analysis from the WGS data. The median sequencing read coverage at the eccDNA/duplication loci was 1.5 fold higher compared to equivalent upstream or downstream regions, suggesting that at least a two fold amplication of one allele occurred in at least 50% of the cells. Surprisingly the eccDNA/duplication events detected by ATAC-seq did NOT show corresponding amplification in WGS (Figure 4B). This result suggests that as with eccDNAs detected by rolling circle amplification, the eccDNA/duplication events identified by ATAC-seq are somatically mosaic in the GBM cell lines and are detected even before a CNV is apparent in the study of a large population of tumor cells.

Finally we analyzed 11 LGG and 8 GBM ATAC-seq libraries and found a total of 423 and 403 eccDNA/duplication events in LGG and GBM samples, respectively. The length distribution of eccDNA/duplications is shown in Figure 4C. 60% of the loci are <1KB (similar to microDNA), but nearly 25% (201 eccDNAs in GBM+LGG) are 50KB to 50MB in length, suggesting that they harbor full length genes. The chromosomal distribution of eccDNA identified in LGG and GBM samples is shown in Figure 4D. The EGFR locus was contained in the eccDNA/duplication identified in GBM patients, supporting our hypothesis that the use of Circle_finder in ATAC-seq data can identify loci that have been amplified even in a subset of the cells in the tumor.

#### Cumulative analysis of all small eccDNA (microDNA)

After pooling all the eccDNA identified so far in this paper, we focused on the ones <1KB to compare their properties with the microDNA we have identified earlier (Kumar et al., 2017; Shibata et al., 2012). We found 762 eccDNA less than 1000 bp. The length distribution of these circles reveals characteristic peaks at ~180 and~380 bases (Figure 5A), that we have noted earlier. The higher GC content relative to the genome average (Figure 5B), and the enrichment of the microDNA from upstream of genes, 5’UTR and CpG islands (Figure 5C) is also similar to our previous reports. Finally around 20% of the small eccDNAs reported here appear to have used flanking sequences of 2-15 base micro homology (Figure 5D) to promote the ligation that gives rise to the circle.

**Figure 5:**
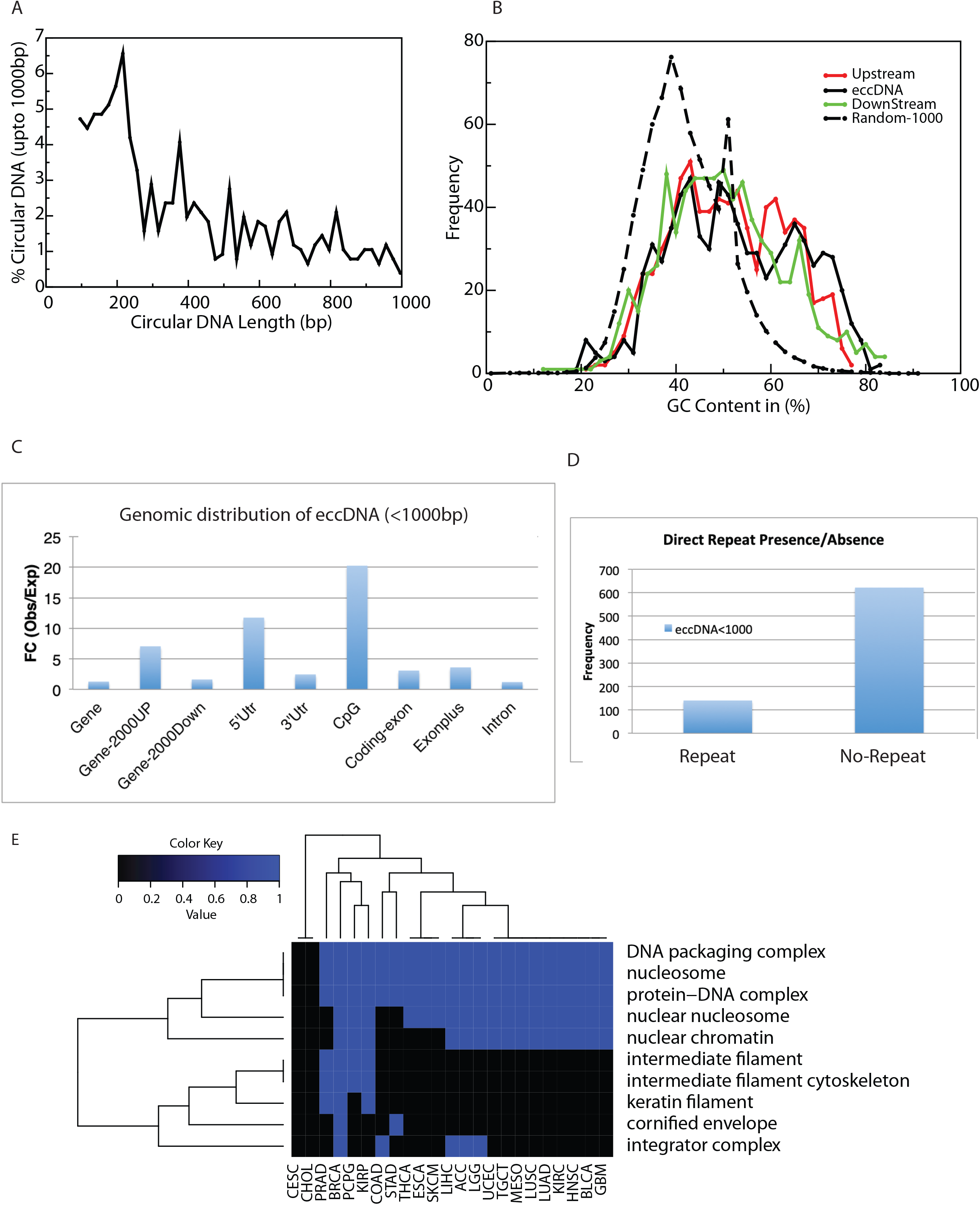
Properties of microDNA identified in this paper (eccDNA<1KB) by ATAC-seq. **(A)** length distribution of eccDNA shows peaks at 180 and 380 bases. **(B)** GC content of eccDNA locus and regions immediately upstream and downstream from the eccDNA is higher than genomic average as calculated from 1000 random stretches of the genome of equivalent length as the eccDNA (Random-1000), (**C)** The sites in the genome that give rise to small eccDNA are enriched relative to random expectation in genic sites, sequences 2 kb upstream from genes and in CpG islands. **(D)** Direct repeats of 2-15 bp flanking the genomic locus of the eccDNA at ligation point are present for ~20% of the loci. **(E)** Gene classes enriched in the set of genes found on the circular DNAs in two or more cancers. If the genes found on the eccDNA/duplication loci in a cancer type are significantly enriched in the indicated pathways, the color in the cell is blue. If the set of genes is not enriched in that cancer, the cell is black.

### Pan cancer analysis of eccDNA in TCGA ATAC-seq data

Finally we analyzed 360 ATAC-seq libraries from twenty-three tumor type generated by TCGA consortium (**Supplementary Table 3**) (Corces et al., 2018). We found a total of 18,143 eccDNAs/duplications of which 86% were <1KB. The unique eccDNA intervals were used to extract the full length genes harbored inside the circle. The cancer driver genes amplified as eccDNA/duplication in individual tumor type is given in Table 2. Gene ontology analysis of all the genes carried on the eccDNA/duplication loci show that Pathways related to nucleosomal events are significantly enriched in these loci (Figure 5E)

## Discussion

We demonstrate that the application of Circle_Finder to ATAC-seq data can identify eccDNA in cell lines and tissues. Most of the eccDNA thus identified in the cell line could be detected by inverse PCR on DNA enriched for extrachromosomal DNA with disenrichment of linear DNA fragments. The metaphase spreads from OVCAR8 cells showed the presence of these eccDNA loci as a signal off the chromosomes, consistent with the loci being extrachromosomal. Even if ATAC-seq is performed without an experimentally dis-enriching linear chromosomal DNA and/or enriching circular DNA, this approach is useful to identify loci that are either contained in eccDNAs or have suffered a tandem segmental duplication in the chromosome. The identification of eccDNA/duplication in the EGFR locus in GBM cell lines as well as GBMs suggested that existing ATAC-seq data from other cancers should also be examined closely to find the driver gene amplification on eccDNA/duplication events in each tumor type. Indeed, we find several cancer driver genes located in such loci (Table 2). These results suggest that deeper sequencing of tumors by ATAC-seq with longer reads will identify many more clinically important sites involved in eccDNA/duplication in these tumors.

Chromosome ends are protected by telomeres. Once the chromosome suffers a catastrophic fragmentation, as in chromothripsis, some parts of the chromosome may be protected from degradation by eccDNA formation. EccDNA can also be generated from extra linear DNA produced by some kind of copying mechanism as a byproduct of DNA replication or repair. Either way, our result suggest that eccDNA are much more prevalent in cancer cell lines and tumors, and that ATAC-seq is an easy method to identify such eccDNA.

It has been reported that longer eccDNA (few kb) may have origin of replication and may get amplified independent of the main chromosome. Thus, if an eccDNA harbors an oncogene, then amplification of such eccDNA in tumor cells will increase the fitness of the tumor cell. In addition, since a centromere is absent in the eccDNA (Turner et al., 2017), eccDNA may segregate unevenly between daughter cells and result in tumor heterogeneity (deCarvalho et al., 2018). Both these mechanisms will increase the likelihood that if a particular type of therapy inhibits a gene resident on a pre-existing eccDNA, then the tumor is likely to acquire resistance through the selective amplification of that eccDNA.

In this context it is particularly exciting that circle (or gene-duplication) at an important locus in a subset of the tumor cells is identified by ATAC-seq even before the amplification is apparent by a CNV analysis of the whole tumor (Fig. 4B). To estimate whether ATAC-seq can identify loci in early (somatically mosaic) states of amplification as eccDNA/segmental duplication, we analyzed the amplicons identified by TCGA from gene array hybridization. It is apparent that an amplicon has to be at least around 1.5 MB long to be detected as a single copy amplification by gene microarrays (3 copies per cell) (Supplementary methods and **Supplementary Fig. 2**). In contrast, we could detect somatically mosaic increase in copy number of loci far smaller than that length. For example, ATAC-seq identifies sites of incipient amplification of the EGFR gene in a subset of tumor cells even before such amplification is detected by copy number measurements, predicting that even if the tumor responds to anti-EGF therapy, it is likely to recur because of amplification of the EGFR gene.

Many of the abundant eccDNA loci intersect with unprocessed pseudogenes, which are known to have introns and regulatory sequences, but crippled by stop codons in the open reading frames (Tutar, 2012). Since eccDNA evolve and pick up substitution, insertion and deletion mutations (Turner et al., 2017; Xu et al., 2019), it is tempting to speculate that amplification of unprocessed pseudogenes on eccDNA and their evolution may make these genes translationally competent to give an unknown advantage during tumorigenesis.

Finally, we note that a large fraction of eccDNA identified by ATAC-seq have properties similar to the microDNA that we reported earlier: length <1KB, with peaks at 180 and 380 bases, high GC content, enrichment of their sites of origin in regions upstream of genes and in CpG islands and the presence of short sequences of homology flanking the chromosomal locus giving rise to the circle. The small size of the circles has thus been confirmed by rolling circle amplification (Shibata et al., 2012), electron microscopy (Shibata et al., 2012) and now by ATAC-seq, ruling out any possibility that the previously reported small size was due to preferential amplification of small circles. The consistent properties of these circles suggest that common mechanisms are involved in their generation in cell lines and in tumors, although it is unclear whether exactly the same mechanisms are involved in producing the longer circles that give rise to gene amplifications.

## Method

### ATAC-seq library preparation

ATAC-seq for cell lines was performed as per the OMNI-ATAC-seq protocol (Corces et al., 2017). Briefly, C4-2 and OVCAR8 cells were grown in RPMI-1640 (Corning #10-040) supplemented with 10% FBS to ~80% confluence. 50,000 viable cells were lysed in 10 mM Tris-HCl (pH 7.4), 10 mM NaCl, 3 mM MgCl_2_ and 0.1% Tween-20. Nuclear pellet was then subjected to transposition reaction using Nextera DNA Sample Preparation kit (Illumina #FC-121-1030) in the presence of 0.01% digitonin and 0.1% Tween-20 at 37°C for 30 minutes and cleaned up with DNA Clean and Concentrator-5 Kit (Zymo #D4014). For qPCR, 3 to 6 additional cycles of PCR amplification was performed using NEBNext High-Fidelity 2X PCR Master Mix (NEB #M0541L) and Nextera Index Kit (Illumina #15055289). Cleaned up libraries were quantified and pooled for sequencing by Novogene.

### Identification of eccDNA from ATAC-seq and WGS libraries

Paired end reads were mapped to the hg38 genome build using bwa-mem (Li and Durbin, 2009) with default setting. The split reads (reads not mapped in contiguous manner) were collected using tool samblaster (Faust and Hall, 2014). If one tag of a paired read is mapped contiguously (one entry in mapped file) and other tag is mapped in a split manner (two entries in mapped file) then the particular read id will have three entries in alignment file. We therefore collected all the read pair IDs that mapped to three unique sites in the genome from the alignment file. Next we collected the split reads that mapped uniquely at two positions on the same chromosome and in the same orientation. Returning to the list of paired end IDs that mapped uniquely to three sites in the genome, we identified paired end IDs where the contiguously mapped read is between the two split reads and on the opposite strand. From this list we annotate a circle if we find at least two junctional sequences (two different pair end reads).

### Copy number amplification (CNA) Analysis

For each identified eccDNA an upstream and downstream genomic interval of equivalent length was created. Next we counted the number of reads that mapped to each of the three intervals (upstream, eccDNA and downstream). Finally CNA was computed by counting the number of mapped read in eccDNA interval divided by mean of the number of reads in upstream and downstream intervals. CNA value more than 1 would suggest the amplification of the locus defined by the eccDNA.

### EccDNA isolation

EccDNA for Figure 3 was prepared from human cancer cell lines as described in Shibata et al. (Shibata et al., 2012). DNA was isolated (HiSpeed Plasmid MidiKit, Qiagen, 12643) according to manufacturer instructions. The DNA, eluted into 1 mL TE was then precipitated by addition of 2 mL of ETOH and 1 ug of glycogen. The DNA was then re-suspended into 20 uL of TE. The linear DNA was then digested with RNaseA and ATP-dependent DNase (Epicentre, E3101K) as per manufacturer instructions. The DNA was then purified with a QIAquick PCR Purification Kit (Qiagen, 28104) to remove salts, enzymes, and nucleotides from the DNA solution.

### Outward directed PCRs (inverse PCR) for detection of eccDNA

Outward directed primers were designed across the junctional tags identified from ATAC-seq analysis. PCR was done with Phusion High-Fidelity DNA polymerase (NEB) according to manufacturers instructions. 3 ng of purified circular DNA was used as template.

Unless otherwise stated, all the computation and plots were made of eccDNA present on chr1-22, chrX & chrY.

### Metaphase FISH

#### Fluorescence in-situ hybridization (FISH)

OVCAR8 cells were cultured in RPMI medium supplemented with 10% FBS and 1% penicillin/streptomycin in presence of 5% C02 in humidified incubator at 370C. Cells were treated with 2 mM thymidine for 16 hours and released for 9 hours in regular medium followed by another block with 2 mM thymidine to arrest the cells at G1/S boundary. The cells were released from the double-thymidine block for 3 hours in regular medium and 9 hours in 0.1 µg/ml Colcemid. Mitotic cells were shaken off, washed twice with 1X PBS and resuspended in 75 mM KCl for 30 min at 370C. The cells were centrifuged at 300Xg for 5 min, fixed with Carnoy’s fixative (3:1 methanol:glacial acetic acid, v/v) on ice for 30 min, washed twice with fixative and metaphase spreads were prepared.

The glass slides containing metaphase spreads were immersed in pre-warmed denaturation buffer (70% formamide, 2X SSC, pH 7.0) at 730C for 5 min and slides were serially dehydrated with ethanol (70%, 85%, 100%) for 2 min each and dried at room temperature until all the ethanol evaporated. The FISH probes (Empire Genomics) were denatured with hybridization buffer at 730C for 5 min and immediately chilled on ice for 2 min. The probe mixture was added onto the slide and coverslips were applied onto the slide and sealed with rubber cement and incubated at 370C for overnight in humidified chamber. The coverslips were removed and slides were washed with pre-warmed 0.4X SSC containing 0.3% NP-40 at 730C for 2 min followed by washing with 2X SCC buffer containing 0.1% NP-40 at room temperature for 5 min. The slides were dried at room and mounted with Vectashield DAPI medium.

### List of TCGA ID that was used for LGG and GBM data analysis

LGG: TCGA-P5-A77X-01A, TCGA-DU-5870-02A, TCGA-DB-A75K-01A, TCGA-W9-A837-01A, TCGA-P5-A72X-01A, TCGA-F6-A8O3-01A, TCGA-FG-A4MY-01A, TCGA-E1-A7YI-01A, TCGA-P5-A735-01A, TCGA-DU-6407-02B.

GBM: TCGA-06-A7TK-01A, TCGA-4W-AA9S-01A, TCGA-OX-A56R-01A, TCGA-76-6656-01A, TCGA-RR-A6KB-01A, TCGA-06-A6S1-01A, TCGA-06-A5U0-01A, TCGA-06-A7TL-01A.

## Acknowledgment

Authors would like to thank dutta lab members for thoughtful discussion on this paper. We thank High Performance Computing Team at the University of Virginia for providing all the support with computation. Thanks to dbGAP & The Cancer Genome Atlas (TCGA) data management teams for data access. Authors also would like to thanks to the patients for their participation in TCGA. Thank to Ryan Corces from Howard Y. Chang group (Howard Hughes Medical Institute, Stanford University, Stanford) for alerting us about raw data availability through TCGA. This work was supported by grants from NIH R01 CA60499, Owens Foundation to Anindya Dutta and a fellowship to SK from the UVA Cancer Center.

## References

Buenrostro, J. D., Wu, B., Chang, H. Y., and Greenleaf, W. J. (2015). ATAC-seq: A Method for Assaying Chromatin Accessibility Genome-Wide. Curr Protoc Mol Biol 109, 21 29 21–29.

Corces, M. R., Granja, J. M., Shams, S., Louie, B. H., Seoane, J. A., Zhou, W., Silva, T. C., Groeneveld, C., Wong, C. K., Cho, S. W., et al. (2018). The chromatin accessibility landscape of primary human cancers. Science 362.

Corces, M. R., Trevino, A. E., Hamilton, E. G., Greenside, P. G., Sinnott-Armstrong, N. A., Vesuna, S., Satpathy, A. T., Rubin, A. J., Montine, K. S., Wu, B., et al. (2017). An improved ATAC-seq protocol reduces background and enables interrogation of frozen tissues. Nat Methods 14, 959–962.

deCarvalho, A. C., Kim, H., Poisson, L. M., Winn, M. E., Mueller, C., Cherba, D., Koeman, J., Seth, S., Protopopov, A., Felicella, M., et al. (2018). Discordant inheritance of chromosomal and extrachromosomal DNA elements contributes to dynamic disease evolution in glioblastoma. Nat Genet 50, 708–717.

Dillon, L. W., Kumar, P., Shibata, Y., Wang, Y. H., Willcox, S., Griffith, J. D., Pommier, Y., Takeda, S., and Dutta, A. (2015). Production of Extrachromosomal MicroDNAs Is Linked to Mismatch Repair Pathways and Transcriptional Activity. Cell Rep 11, 1749–1759.

Kumar, P., Dillon, L. W., Shibata, Y., Jazaeri, A. A., Jones, D. R., and Dutta, A. (2017). Normal and Cancerous Tissues Release Extrachromosomal Circular DNA (eccDNA) into the Circulation. Mol Cancer Res 15, 1197–1205.

Libermann, T. A., Nusbaum, H. R., Razon, N., Kris, R., Lax, I., Soreq, H., Whittle, N., Waterfield, M. D., Ullrich, A., and Schlessinger, J. (1985). Amplification, enhanced expression and possible rearrangement of EGF receptor gene in primary human brain tumours of glial origin. Nature 313, 144–147.

Maher, C. A., and Wilson, R. K. (2012). Chromothripsis and human disease: piecing together the shattering process. Cell 148, 29–32.

Maire, C. L., and Ligon, K. L. (2014). Molecular pathologic diagnosis of epidermal growth factor receptor. Neuro Oncol 16 Suppl 8, viii1–6.

Moller, H. D., Larsen, C. E., Parsons, L., Hansen, A. J., Regenberg, B., and Mourier, T. (2015a). Formation of Extrachromosomal Circular DNA from Long Terminal Repeats of Retrotransposons in Saccharomyces cerevisiae. G3 (Bethesda) 6, 453–462.

Moller, H. D., Mohiyuddin, M., Prada-Luengo, I., Sailani, M. R., Halling, J. F., Plomgaard, P., Maretty, L., Hansen, A. J., Snyder, M. P., Pilegaard, H., et al. (2018). Circular DNA elements of chromosomal origin are common in healthy human somatic tissue. Nat Commun 9, 1069.

Moller, H. D., Parsons, L., Jorgensen, T. S., Botstein, D., and Regenberg, B. (2015b). Extrachromosomal circular DNA is common in yeast. Proc Natl Acad Sci U S A 112, E3114–3122.

Shibata, Y., Kumar, P., Layer, R., Willcox, S., Gagan, J. R., Griffith, J. D., and Dutta, A. (2012). Extrachromosomal microDNAs and chromosomal microdeletions in normal tissues. Science 336, 82–86.

Shoura, M. J., Gabdank, I., Hansen, L., Merker, J., Gotlib, J., Levene, S. D., and Fire, A. Z. (2017). Intricate and Cell Type-Specific Populations of Endogenous Circular DNA (eccDNA) in Caenorhabditis elegans and Homo sapiens. G3 (Bethesda) 7, 3295–3303.

Turner, K. M., Deshpande, V., Beyter, D., Koga, T., Rusert, J., Lee, C., Li, B., Arden, K., Ren, B., Nathanson, D. A., et al. (2017). Extrachromosomal oncogene amplification drives tumour evolution and genetic heterogeneity. Nature 543, 122–125.

Tutar, Y. (2012). Pseudogenes. Comp Funct Genomics 2012, 424526.

Xie, Q., Wu, T. P., Gimple, R. C., Li, Z., Prager, B. C., Wu, Q., Yu, Y., Wang, P., Wang, Y., Gorkin, D. U., et al. (2018). N(6)-methyladenine DNA Modification in Glioblastoma. Cell 175, 1228–1243 e1220.

Xu, K., Ding, L., Chang, T. C., Shao, Y., Chiang, J., Mulder, H., Wang, S., Shaw, T. I., Wen, J., Hover, L., et al. (2019). Structure and evolution of double minutes in diagnosis and relapse brain tumors. Acta Neuropathol 137, 123–137.

