## Supplementary-Material-Table-Figure for "ATAC-seq identifies thousands of extrachromosomal circular DNA in cancers and cell lines"

**Supplementary Material Method:**

We tested whether the detection of eccDNAs from ATAC-Seq data can identify somatically mosaic amplifications before they can be detected by copy number variation analyses from genotyping array data. To determine the sensitivity of detection of an amplicon by genotyping arrays, we downloaded the previously released copy number variation (CNV) results generated by the TCGA research network. The algorithm used by the TCGA research network segments the chromosomes into smaller sections where an amplification or deletion is detected. Empirically, the resulting length of segments with CNV determined by the algorithm are the result of (1) the true length of the amplified or deleted segment and (2) the extent to which the segment was amplified or deleted. While we cannot know whether or not a reported CNV-segment should have been further segmented, we hypothesized that if we analyzed ten segments with a similar level of amplification, the smallest length among them approximates the smallest length that can be detected by the algorithm at that level of amplification, since the power to detect CNV changes increases with as the extent of amplification increases

The TCGA research network reported amplifications as segment mean >0 , where segment mean is ln(Copy number/2). All segments with segment mean >0.1 were ordered by reported segment mean values. Bins of ten segments were analyzed for the smallest segment in each bin. The median segment mean value of each bin (extent of amplification) is plotted against and the log-transformed smallest segment length in that bin (Figure X).

The correlation between the segment length and segment amplification can be modeled as a linear function with the following formula:

ln(Minimum Segment Length) = 15.8304 – 2.7475*Median Segment Mean

This relatively simple model accurately captured the relationship between minimum segment length and the segment mean (Adjusted R2 = 0.5442, p<2.2E-16).

If one extra copy of an amplicon is present in every single cell of the sample, the segment mean value is 0.585 [log2(3/2)]. From the linear model in Figure X, the minimum segment length detectable at this segment mean value is 1.5 MB. Therefore most of the somatically mosaic amplifications driven by most of the eccDNAs in our study (median length~ 2 KB) will not be captured using genotyping arrays.

**Supplementary Figure1:** eccDNA in GBM cell lines. **(A)** Length distribution of identified eccDNA in GBM cell lines. **(B)** Karyotype plot showing chromosomal distribution of eccDNA identified in GBM cell lines.

**Supplementary Figure 2:** In general, genotyping arrays will fail to identify amplifications caused by the circular DNAs detected in our analysis. The minimum detectable segment length (y-axis) is partially dependent on the extent of amplification (x-axis). Assuming that one copy of a circular DNA is present in every cell of a patient's tumor, the segment mean would be 0.585 and the minimum detectable length would therefore be approximately 1.5 MB. Given that the average length of the circular DNAs detected in this analysis was 2KB, the circular DNAs detected in this analysis would not show up as amplifications when genotyping array data is analyzed.

**Supplementary Table 1:** Common eccDNA between WGS & ATACseq libraries in TCGA-DU-5870-02A

| | JT-ATAC | JT-WGS | Length |
|---|---|---|---|
| chr11:109186224-109186306 | 1 | 2 | 82 |
| chr12:8208854-8238302 | 1 | 10 | 29448 |
| chr14:56200142-56200312 | 2 | 3 | 170 |
| chr14:61172784-61172905 | 1 | 3 | 121 |
| chr14:99811139-99811333 | 1 | 2 | 194 |
| chr15:101390179-101390341 | 1 | 3 | 162 |
| chr15:52523965-52524853 | 1 | 6 | 888 |
| chr16:3142266-3142362 | 2 | 1 | 96 |
| chr16:69820782-69825118 | 2 | 10 | 4336 |
| chr17:80024303-80024653 | 1 | 9 | 350 |
| chr19:49867632-49867701 | 1 | 2 | 69 |
| chr20:58695018-58695339 | 1 | 7 | 321 |
| chr22:49226585-49228448 | 1 | 11 | 1863 |
| chr2:110101070-110104430 | 1 | 6 | 3360 |
| chr3:120222954-120223088 | 1 | 1 | 134 |
| chr3:42898790-46004380 | 1 | 1 | 3105590 |
| chr4:150242941-150244988 | 4 | 11 | 2047 |
| chr4:156660547-156660686 | 2 | 2 | 139 |
| chr4:187673040-187673173 | 1 | 3 | 133 |
| chr5:134246913-134247079 | 1 | 3 | 166 |
| chr6:120504095-120504276 | 1 | 2 | 181 |
| chr6:167516501-167516632 | 1 | 2 | 131 |
| chr7:65038315-65873352 | 3 | 2 | 835037 |
| chrX:49069208-49069329 | 2 | 1 | 121 |
| chrY:10850436-56834809 | 1 | 1 | 45984373 |

**Supplementary Table 2:** Common eccDNA between WGS & ATACseq libraries in TCGA-DU-5870-02A

| | JT-ATAC | JT-WGS | Length |
|---|---|---|---|
| chr10:54142247-55002600 | 2 | 1 | 860353 |
| chr10:95447028-95448268 | 1 | 2 | 1240 |
| chr11:28824878-36215494 | 1 | 10 | 7390616 |
| chr11:29133708-29430833 | 1 | 2 | 297125 |
| chr11:31877072-36188811 | 3 | 8 | 4311739 |
| chr11:36202059-36238115 | 2 | 12 | 36056 |
| chr11:36214300-36237662 | 1 | 5 | 23362 |
| chr11:36412110-36417948 | 1 | 4 | 5838 |
| chr13:101820637-101820755 | 1 | 1 | 118 |
| chr13:76789088-76789267 | 1 | 1 | 179 |
| chr15:91440230-91446387 | 1 | 11 | 6157 |
| chr18:56826056-56826276 | 1 | 3 | 220 |
| chr2:11554-91086 | 1 | 6 | 79532 |
| chr2:16052063-16452027 | 1 | 3 | 399964 |
| chr2:16177696-17516120 | 1 | 97 | 1338424 |
| chr2:16225124-16226720 | 2 | 89 | 1596 |
| chr2:16567034-17412029 | 1 | 1 | 844995 |
| chr2:17279980-17334701 | 4 | 10 | 54721 |
| chr2:17296668-17297551 | 2 | 11 | 883 |
| chr2:17584356-17877776 | 2 | 13 | 293420 |
| chr4:52081945-52082092 | 1 | 3 | 147 |
| chr6:118690555-118692765 | 1 | 4 | 2210 |
| chr6:54059859-54063911 | 1 | 7 | 4052 |
| chr7:116674913-121917354 | 5 | 25 | 5242441 |
| chr8:145036620-145051835 | 1 | 10 | 15215 |
| chrY:10945178-11295108 | 2 | 12 | 349930 |

**Supplementary Table 3**: Number of ATAC-seq libraries analyzed for each tumor type

| | |
|---|---|
| ACCx | 8 |
| BLCA | 10 |
| BRCA | 70 |
| CESC | 3 |
| CHOL | 1 |
| COAD | 38 |

| | |
|---|---|
| ESCA | 17 |
| GBMx | 8 |
| HNSC | 9 |
| KIRC | 15 |
| KIRP | 29 |
| LGGx | 11 |
| LIHC | 15 |
| LUAD | 21 |
| LUSC | 12 |
| MESO | 5 |
| PCPG | 9 |
| PRAD | 21 |
| SKCM | 9 |
| STAD | 19 |
| TGCT | 8 |
| THCA | 12 |
| UCEC | 10 |

A

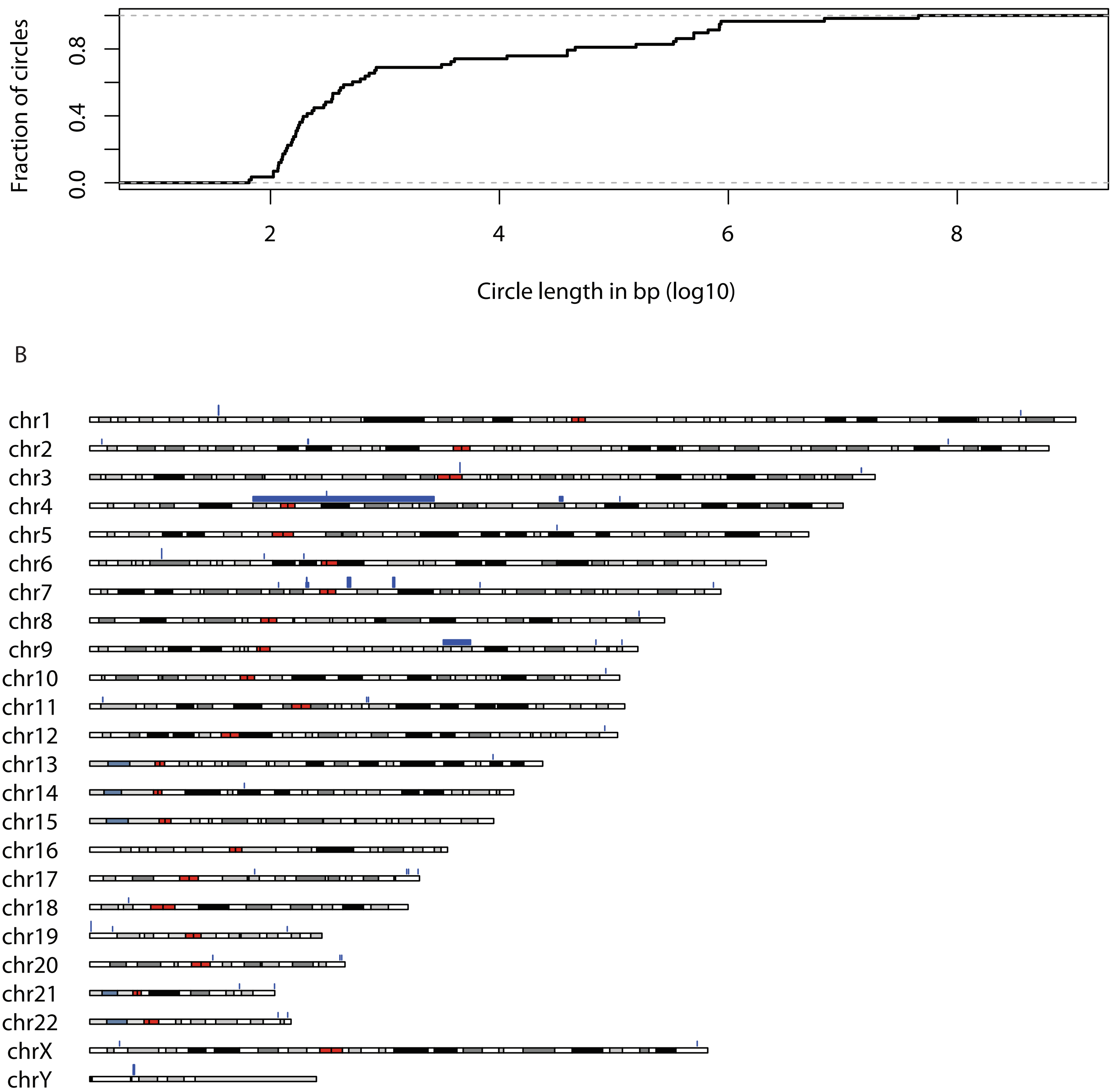


eccDNA distribution on various chromososmes (GBM cell lines)

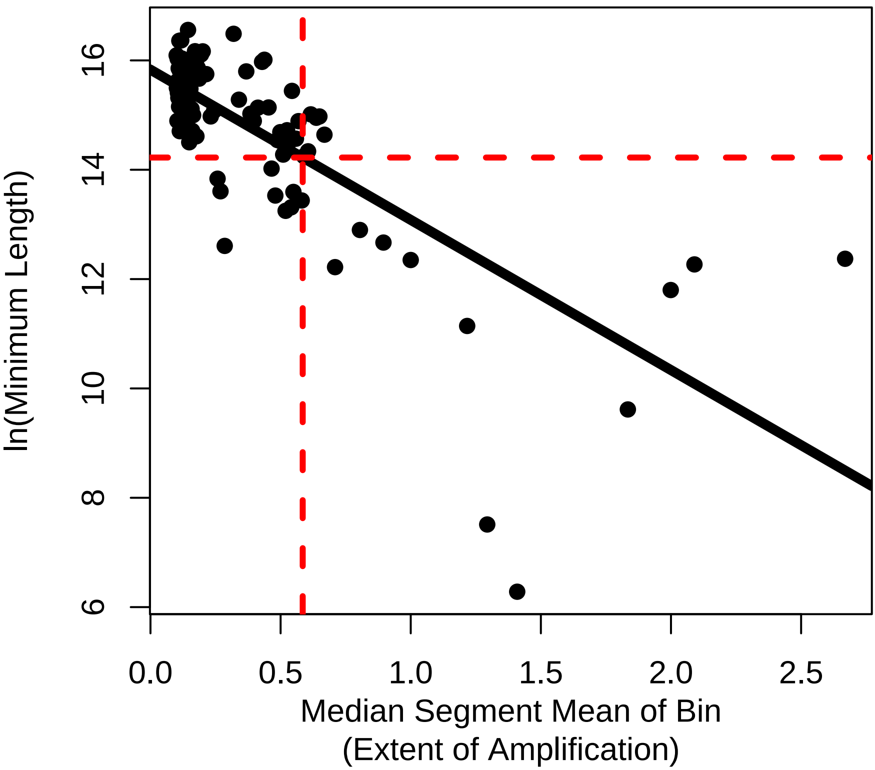

16
14
12
10
8
6
ln(Minimum Length)
0.0
0.5
1.0
1.5
2.0
2.5
Median Segment Mean of Bin
(Extent of Amplification)
